# ACD15, ACD21 and SLN regulate accumulation and mobility of MBD6 to silence genes and transposable elements

**DOI:** 10.1101/2023.08.23.554494

**Authors:** Brandon A. Boone, Lucia Ichino, Shuya Wang, Jason Gardiner, Jaewon Yun, Yasaman Jami-Alahmadi, Jihui Sha, Cristy P. Mendoza, Bailey J. Steelman, Aliya van Aardenne, Sophia Kira-Lucas, Isabelle Trentchev, James A. Wohlschlegel, Steven E. Jacobsen

**Author notes:** These authors contributed equally (Brandon A. Boone, Lucia Ichino). Corresponding author. (S.E.J).

## Abstract

DNA methylation mediates silencing of transposable elements and genes in part via recruitment of the Arabidopsis MBD5/6 complex, which contains the methyl-CpG-binding domain (MBD) proteins MBD5 and MBD6, and the J-domain containing protein SILENZIO (SLN). Here we characterize two additional complex members: α-crystalline domain containing proteins ACD15 and ACD21. We show that they are necessary for gene silencing, bridge SLN to the complex, and promote higher order multimerization of MBD5/6 complexes within heterochromatin. These complexes are also highly dynamic, with the mobility of complex components regulated by the activity of SLN. Using a dCas9 system, we demonstrate that tethering the ACDs to an ectopic site outside of heterochromatin can drive massive accumulation of MBD5/6 complexes into large nuclear bodies. These results demonstrate that ACD15 and ACD21 are critical components of gene silencing complexes that act to drive the formation of higher order, dynamic assemblies.

**One-Sentence Summary:** Arabidopsis ACD21 and ACD15 drive accumulation of MBD5/6 complex silencing assemblies at methyl-CG sites and recruit SLN to maintain protein mobility in these assemblages.

## Main Text

Eukaryotic organisms must properly localize macromolecules within cells to maintain homeostasis. Membrane bound organelles serve this purpose, but recent discoveries have revealed the existence of membrane-less organelles or compartments(*1*–*3*). Often referred to as biological condensates, liquid-liquid phase separated (LLPS) condensates, or supramolecular assemblies, these compartments such as stress granules, p-granules, heterochromatin, and the nucleolus concentrate proteins and nucleic acids to facilitate specific and efficient processes(*4*–*7*). If not properly controlled, the accumulation of these protein assemblies can lead to aggregates with detrimental impacts on cellular homeostasis and disease, yet how cells regulate these assemblies remains unclear(*3, 8, 9*). Molecular chaperones, such as heat shock proteins (HSPs), serve highly conserved roles to regulate the solubility, folding, and aggregation of proteins within cells, making them obvious candidates for the regulation of biological condensates(*10*–*14*). Small HSPs (sHSPs) use their conserved α-crystalline domains (ACD) to form dimers which then create large and dynamic oligomeric assemblies that act as first line of defense against protein aggregation via a “holdase” activity(*14*). sHSPs further recruit J-domain containing proteins (JDPs) which act as co-chaperones for HSP70 proteins to maintain protein homeostasis (*14*–*17*). Both sHSPs and JDP/HSP70 pairs have been shown to associate with and regulate disease related cellular condensates across species(*18*–*21*).

In *Arabidopsis thaliana*, pericentromeric heterochromatin is organized into compartments called chromocenters that are chromatin dense regions containing most of the DNA methylated and constitutively silenced TEs and genes, as well as heterochromatic proteins such as DNA methylation binding complexes (*13, 22*–*26*). Previous work has shown that multiple Arabidopsis DNA methylation binding complexes silence or promote expression of genes through recruitment of molecular chaperones with unknown functions (*27*–*29*). For example, MBD5 and MBD6 redundantly silence a subset of TEs and promoter-methylated genes via recruitment of SLN, a JDP. MBD5/6 also interact with two ACD containing proteins called ACD15.5/RDS2 and ACD21.4/RDS1, hereafter referred to as ACD15 and ACD21(*29*–*31*). While ACD15 and ACD21 have been implicated in silencing of a transgene reporter, their specific chromatin functions remain unknown (*31*). Here we demonstrate that ACD15 and ACD21 are necessary and sufficient for the accumulation of high density MBD6 at methylated CG sites to silence genes and TEs, while also bridging SLN to MBD5/6 to maintain the high mobility of all complex components. We further demonstrate that MBD5/6 complex assemblies can be formed at discrete foci outside of chromocenters, in an ACD15 and ACD21 dependent manner, to cause gene silencing.

### ACD15 and ACD21 colocalize with MBD5 and MBD6 genome-wide and are essential for silencing

We previously observed that MBD5, MBD6 and SLN pulled-down two ACD containing proteins named ACD15 and ACD21 (*29*). To investigate their binding patterns on chromatin, we performed Chromatin Immunoprecipitation sequencing (ChIP-seq) of FLAG-tagged ACD15 and ACD21. We observed that all five proteins colocalized genome-wide, and none of them appeared to have truly unique ChIP-seq peaks, suggesting that they could be recruited to DNA together as a complex (Fig. 1A-B, S1A). Furthermore, ACD15 and ACD21 showed a non-linear correlation with meCG density similar to MBD6 and SLN(*29*) suggesting that MBD5/6 complex members all accumulate preferentially at high density meCG sites (Fig. 1C).

**Fig. 1.**
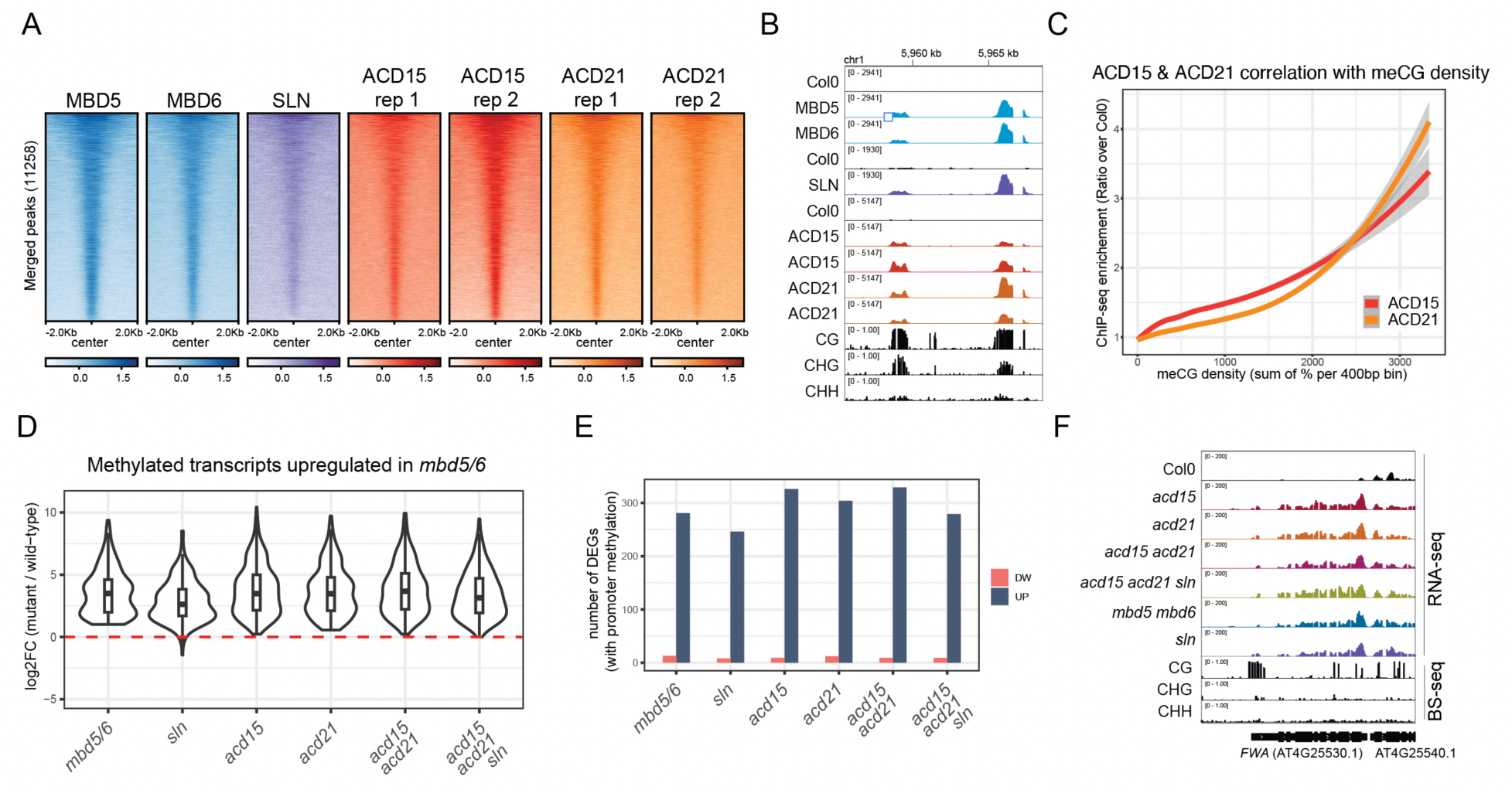
ACD15 and ACD21 are required for silencing. **(A)** Heatmap of FLAG-tagged MBD5, MBD6, SLN, ACD15, and ACD21 ChIP-seq enrichment (log2FC over no-FLAG Col0 control) centered at all merged peaks. **(B)** Genome browser image of ChIP-seq data showing two methylated loci co-bound by all MBD5/6 complex members. **(C)** Loess curves showing correlation between ChIP-seq enrichment for a representative replicate and CG methylation density. **(D)** Violin plots showing mature pollen RNA-seq data for the indicated mutants, at *mbd5 mbd6* upregulated transcripts (6 replicates per genotype). **(E)** Comparison between genotypes of the number of RNA-seq differentially expressed genes (DEGs) with >40% CG methylation levels around the TSS. **(F)** Genome browser image of RNA-seq data at the *FWA* locus in the indicated genotypes. Wild-type BS-seq data is shown as reference.

To test whether ACD15 and ACD21 are required for silencing we generated *acd15* and *acd21* single mutants, an *acd15 acd21* double mutant, and an *acd15 acd21 sln* triple mutant via CRISPR/Cas9 (fig. S1B). RNA-seq analysis revealed that all mutants showed very similar transcriptional derepression patterns at DNA methylated genes and TEs as compared to *mbd5 mbd6* and *sln* mutants (Fig. 1D-F, S1C-D). This includes the *FWA* gene which was previously shown to be silenced by the MBD5/6 complex (Fig. 1F) (*29*). These results demonstrate that ACD15 and ACD21 are critical components of the MBD5/6 complex required for silencing.

### ACD15 and ACD21 bridge SLN to MBD5 and MBD6

To determine the specific organization of the MBD5/6 complex, we performed IP-MS experiments using FLAG-tagged transgenic lines for each complex component in different mutant backgrounds (Fig. 2A, Table S1). In the wild type Col0 background, ACD15 and ACD21 pulled-down each other, MBD5, MBD6, SLN, and the same HSP70 proteins that were found to interact with the MBD5/6 complex previously (*29*) (Fig. 2A, Table S1). MBD5 and MBD6 pulled down peptides of ACD15 and ACD21 in the absence of SLN, while SLN did not pull down MBD5 and MBD6 in the absence of ACD15 and ACD21, suggesting that ACD15 and ACD21 bridge the interaction between MBD5/6 and SLN (Fig. 2A-B). Consistent with this model, ACD15 and ACD21 pulled down MBD5 and MBD6 in the *sln* mutant background, and SLN pulled down ACD15 and ACD21 in the *mbd5 mbd6* mutant background (Fig. 2A). ACD15 also pulled-down MBD5 and MBD6 but not SLN in the absence of ACD21, while ACD21 did not pull down MBD5 and MBD6 in the absence of ACD15 (Fig. 2A). These results suggest that the MBD5/6 complex is organized such that MBD5 or MBD6 interact with ACD15, ACD15 interacts with ACD21, and ACD21 interacts with SLN (*29*) (Fig. 2B).

**Fig. 2.**
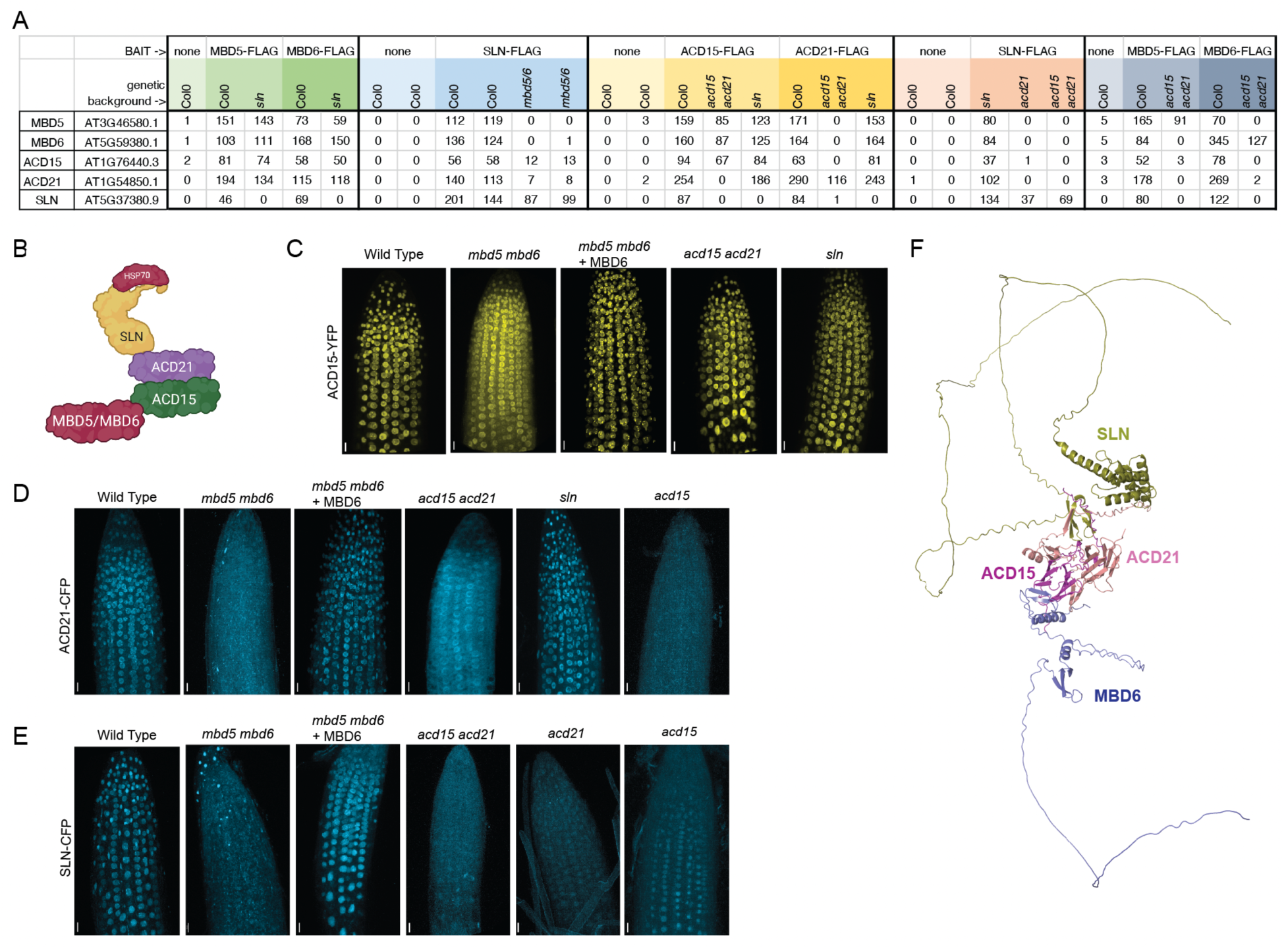
ACD15 and ACD21 bridge SLN to MBD5/6. **(A)** IP-MS of flag-tagged MBD5/6 complex members in the indicated genetic backgrounds (MS/MS counts). **(B)** MBD5/6 complex organization as predicted by IP-MS. Created with BioRender.com (**C-E)** 3D reconstruction of root meristems of plants expressing fluorescently tagged ACD15, ACD21, or SLN in wild-type (Col0) and mutant backgrounds. Scale bar = 20 μm. **(F)** Predicted structure of MBD5/6 complex from AlphaFold Multimer (*33*). MBD6=Blue, ACD15=magenta, ACD21=maroon, SLN=gold.

To further study the organization and localization of MBD5/6 complex components we used live confocal imaging of root tips to determine the cellular localization of fluorescent-protein-tagged ACD15, ACD21, SLN, and MBD6. In wild-type plants, ACD15, ACD21, and SLN all showed clear nuclear localization which correlated strongly with nuclear MBD6 (Fig. 2C-E and S2A-C). ACD21, ACD15, and SLN all showed an increase in cytosolic signal in *mbd5 mbd6* mutant plants which was rescued by coexpressing MBD6, demonstrating that all members of the complex require genetically redundant MBD5 or MBD6 for proper nuclear localization (Fig. 2C-E). The reduction of nuclear localization of SLN is also consistent with previous ChIP-seq experiments showing loss of chromatin bound SLN in the absence of MBD5 and MBD6(*29*). ACD15 maintained nuclear localization and correlation with MBD6 in *acd15 acd21* and *sln* mutant plants whereas ACD21 lost nuclear localization and correlation with MBD6 in *acd15* and *acd15 acd21* mutants, but not in the *sln* mutant (Fig. 2C-D, S2D-G). Finally, SLN nuclear localization and correlation with MBD6 decreased in *acd15, acd21*, and *acd15 acd21*, mutant plants (Fig. 2E, S2H,I). These results demonstrate that ACD21 requires ACD15 for proper nuclear localization, while SLN requires both ACD15 and ACD21, consistent with the complex organization model suggested by IP-MS experiments (Fig. 2B).

We used the protein folding algorithm AlphaFold Multimer to predict protein-protein interactions within the MBD5/6 complex(*32, 33*). AlphaFold Multimer confidently predicted that ACD15 interacts with MBD6 (or MBD5), that ACD15 interacts with ACD21, and that ACD21 interacts with SLN, all consistent with our experimental data from IP-MS and confocal microscopy (Fig. 2F). When given two copies of each member of the complex (MBD6, ACD15, ACD21, and SLN), AlphaFold Multimer also confidently predicted that ACD15 and ACD21 form a dimer of two heterodimers in the middle of the structure, suggesting that the MBD5/6 complex likely contains at least two copies of each protein (fig. S2J-L). This is consistent with previous results showing dimer formation by other ACD containing sHSPs (*34*). Given the genetic redundancy of MBD5 and MBD6, the complex would be predicted to contain a minimum of two MBD5s, two MBD6s, or one MBD5 plus one MBD6 (fig. S2L). In line with this prediction, we found that MBD5 and MBD6 pull-down each other in IP-MS data in wild-type, but not in the *acd15 acd21* double mutant background, indicating that ACD15/ACD21 facilitate interaction between two MBD5/6 proteins (Fig. 2A, S2L).

### ACD15, ACD21, and SLN regulate heterochromatic localization, accumulation, and dynamics of the MBD5/6 complex

Given the known role of molecular chaperones in the regulation of protein complexes and aggregates (*35*), we hypothesized that ACD15, ACD21, and SLN may regulate the dynamics of MBD5/6 nuclear complexes. To test this, we measured the nuclear localization and mobility of MBD6 in root cells using live-cell, fluorescence, confocal microscopy. In wild-type and *mbd5 mbd6* mutant plants, MBD6 formed foci, which colocalized with ACD15, ACD21, and SLN foci (Fig. 3A-B, S3A). MBD6 foci also overlapped with DAPI-staining chromocenters, as previously shown when MBD6 was overexpressed in leaf cells (fig. S3B) (*36*). To measure the mobility of MBD6 protein we used fluorescence recovery after photobleaching (FRAP) experiments (*37*). FRAP in wild-type plants revealed that MBD6 moves rapidly within nuclei with a FRAP recovery half time (t_1/2_) of ∼3.60 seconds back into chromocenters after bleaching (Fig. 3C-D, S3D).

**Fig. 3.**
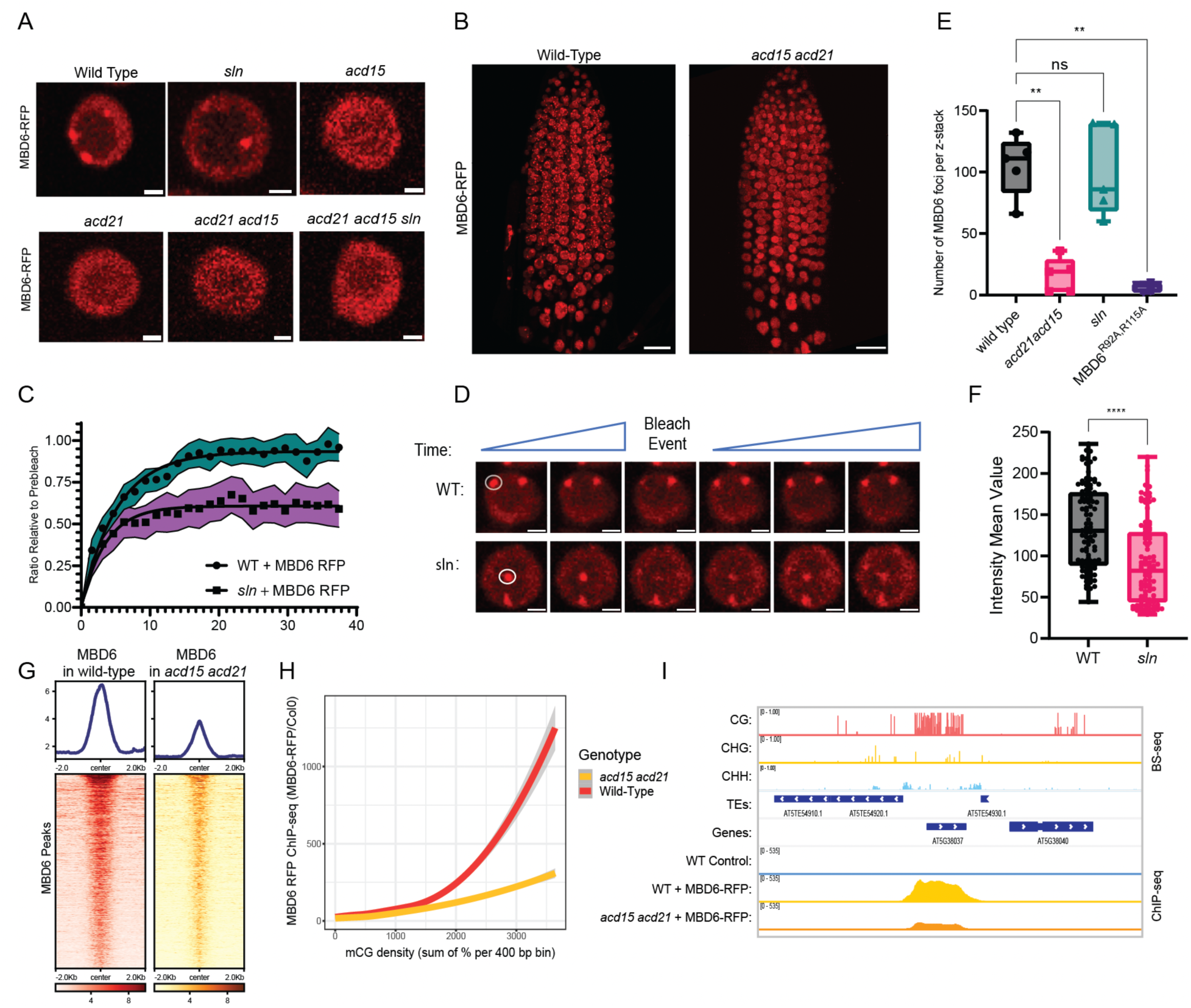
ACD15, ACD21, and SLN regulate MBD6 accumulation and mobility. **(A)** Representative MBD6-RFP nuclear images in mutant backgrounds. Scale bar = 2μM. **(B)** 3D reconstruction of MBD6-RFP root meristem z-stacks. Scale bar = 20μM. **(C)** FRAP recovery curves comparing MBD6 signal in WT and *sln* plants. Shaded area: 95% confidence interval of FRAP data (N=25 from 5 plants lines), dots: mean values, line: fitted one-phase, non-linear regression. **(D)** Representative image of FRAP experiment. White circles indicate foci chosen for bleaching. Scale bars = 2μM **(E)** MBD6 foci counts across 50 slice Z-stacks of root meristems from five plant lines per genotype. Welch’s ANOVA and Dunnet’s T3 multiple comparisons test (**: P<0.01, NS: P>=0.05). **(F)** Box plots of mean intensity values of MBD6 foci (5 individual plants per genotype). Two-tailed t-test (****: P<0.0001). **(G)** Heatmaps and metaplots of MBD6-RFP ChIP-seq signal (log2 ratio over no-FLAG Col0 control) at peaks called in “MBD6-RFP in wild-type” dataset. **(H)** Loess curves showing correlation between MBD6-RFP ChIP-seq enrichment and CG methylation density. **(I)** Genome browser tracks showing an example of a high density meCG site bound by MBD6-RFP (ChIP-Seq). Wild-type BS-seq data is shown as reference.

We next tested whether MBD6 nuclear distribution or mobility was altered in *sln* mutants. Although MBD6 formed a similar number of nuclear foci in *sln* compared to wild-type plants, these foci showed somewhat reduced fluorescence intensity, suggesting that MBD6 was accumulating less efficiently within heterochromatin (Fig. 3A, E, F). FRAP of MBD6 in *sln* mutant plants revealed a dramatic reduction in mobility and a lack of full recovery of signal post bleaching (Fig. 3C-D and S3D). Similar FRAP experiments on ACD15 and ACD21 nuclear foci showed that both were highly mobile in wild-type (t_1/2_ of 3.63 and 4.30 seconds respectively), but were much less mobile and failed to recover full signal in *sln* mutant plants (Figure S3D-H), and also showed decreased fluorescence intensity of foci in *sln* compared to wild-type (fig. S3I-J). SLN thus regulates the mobility, and to a lesser extent the accumulation of the MBD5/6 complex.

Given the IP-MS, microscopy, and structure prediction results showing that ACD15 and ACD21 bridge the interaction between MBD6 and SLN we expected *acd15 acd21* mutants to alter the FRAP mobility of MBD6 in a manner similar to *sln* mutants. However, we found that the number of MBD6 foci were dramatically lower in *acd15 acd21* mutant plants compared to wild-type plants, with only occasional MBD6 foci observed (Fig. 3A, B, E). Instead, MBD6 nuclear signal in *acd15 acd21* mutant plants was more diffusely distributed across nuclei compared to either wild-type or *sln* plants (Fig. 3A). A decreased number of MBD6 foci and a lack of overlap of these foci with DAPI stained chromocenters was also observed in *acd15, acd21*, and *acd15 acd21 sln* mutant plants (Fig. 3A, fig. S3C). Thus, ACD15 and ACD21 are required for MBD6 to efficiently concentrate into nuclear foci. This effect was specific to ACD15 and ACD21 since loss of IDM3 (LIL), an ACD protein in the MBD7 complex (*28*), did not affect the MBD6 nuclear foci (fig. S3K-L).

We performed ChIP-seq on MBD6 in *acd15 acd21* mutant plants to quantify the impact on MBD6 chromatin localization. In wild-type plants, MBD6 localized to previously published MBD6 peaks(*29*) and showed a non-linear correlation with meCG density, displaying strong enrichment at highly dense methylated regions (Fig. 3G-I). However, MBD6 chromatin enrichment in *acd15 acd21* mutant plants, although not abolished, was decreased dramatically and showed much less preference for binding to high density meCG sites (Fig. 3G-I). Thus, while ACD15/ACD21 are not necessary for MBD6 to bind meCG sites, they are needed for high accumulation of MBD6 at high density meCG sites, which is consistent with the decrease of observable MBD6 foci in *acd* mutants (Fig. 3A-B).

Taken together, these results demonstrate that ACD15 and ACD21 are required for high level accumulation of MBD5/6 complexes in chromocenters and at high density meCG sites, while SLN regulates the mobility of these complexes to maintain dynamic recycling of proteins.

### The StkyC domain of MBD6 is required for gene silencing and recruits ACD15 to the complex

The AlphaFold-predicted structure of MBD6 reveals two structured domains, the MBD and a C-terminal domain of unknown function, as well as two intrinsically disordered regions (IDRs) (Figure 4A, and S4A). The C-terminal folded domain shares amino acid similarity with the C-terminus of two related MBD proteins, MBD5 and MBD7 (fig. S4B). This region of MBD7 has been termed the StkyC domain, and is the proposed binding site for the ACD containing IDM3 protein, which belongs to the same family as ACD15 and ACD21(*28*). This suggests that the StkyC of MBD6 would interact with ACD15, and indeed this interaction is confidently predicted by AlphaFold Multimer (Fig. 2F, S2J-K, S4A).

**Fig. 4.**
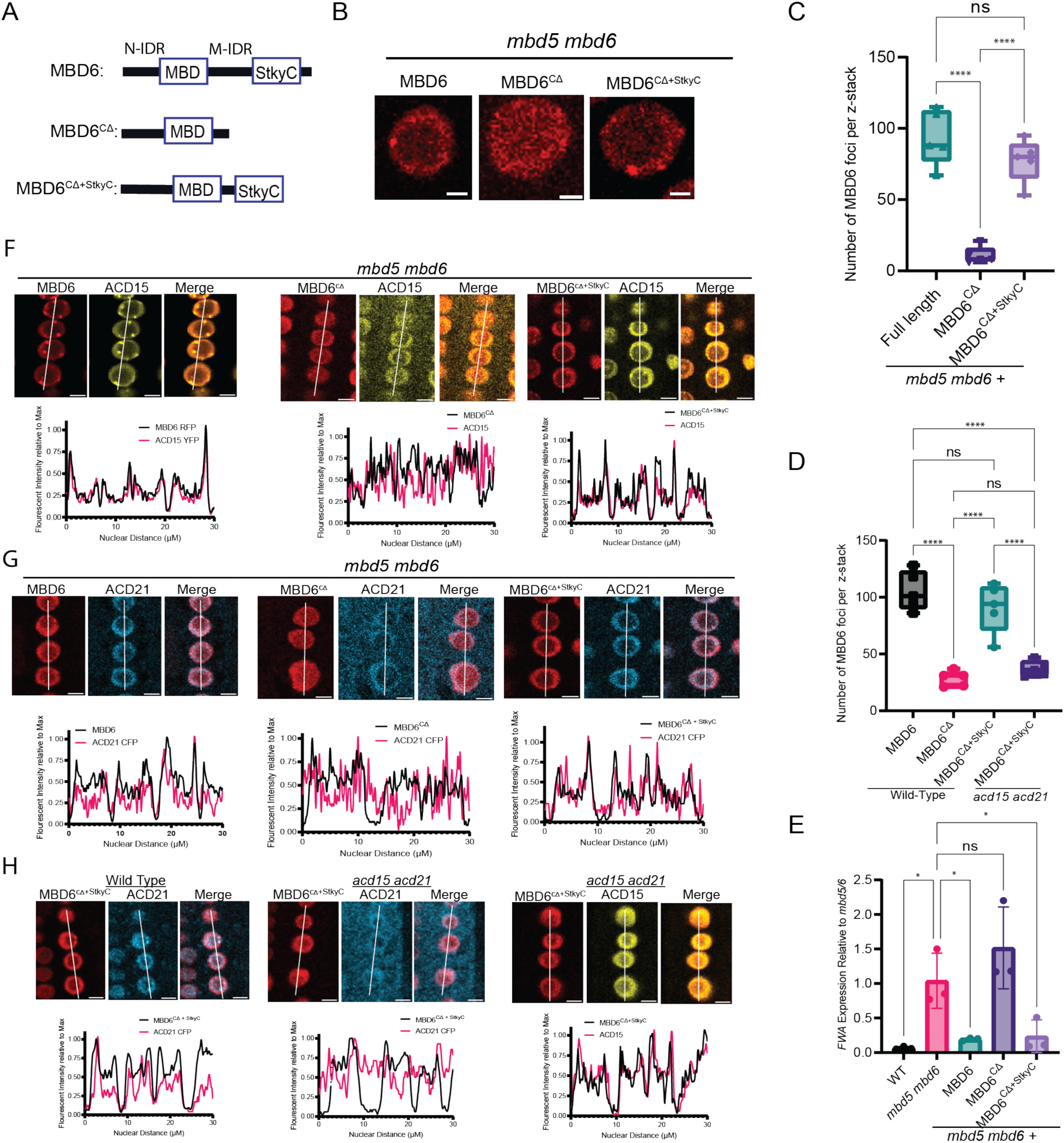
StkyC domain of MBD6 is necessary for function and localization of MBD6. **(A)** Graphical description of MBD6 mutant constructs. **(B)** Representative nuclei showing MBD6-RFP signal. Scale bar = 2 μM. **(C-D)** Number of MBD6 nuclear foci (5 different plants lines per sample, Z-stack of 50 slices). Brown-Forsythe ANOVA with Tukey’s multiple corrections test (****: P<0.0001, NS: P>=0.05). **(E)** *FWA* expression from RT-qPCR of flower bud RNA. Brown-Forsythe ANOVA with Dunnet’s multiple corrections test (*: P<0.05, NS: P>=0.05). **(F-H)** Representative images and nuclear profile plots of MBD6-RFP mutants with either ACD15-YFP or ACD21-CFP. White lines indicate the region plotted in the graphs (“nuclear distance”). Scale bars = 10μM.

To experimentally determine what domains of MBD6 are necessary for gene silencing and chaperone interactions we first truncated the N-terminus (MBD6^NΔ^ (leaving amino acids 66-224)) or the C-terminus of MBD6 (MBD6^CΔ^ (leaving amino acids 1-146)) (fig. S4C). To test if these mutants are functional for silencing, we performed RT-qPCR of the *FWA* gene, a target of the MBD5/6 complex (Fig. 1F)(*29*), in *mbd5 mbd6* mutant plants expressing full-length or truncated MBD6 alleles. *FWA* derepression in *mbd5 mbd6* plants was rescued by full length MBD6-RFP or MBD6^NΔ^-RFP, but not by MBD6^CΔ^-RFP, showing that the middle IDR and/or the StkyC domain are required for MBD6 function (fig. S4D). MBD6^CΔ^ also showed a dramatic reduction in nuclear foci compared to full length MBD6, a phenotype similar to that observed in *acd15 acd21* mutants and consistent with loss of the ACD15 binding site (Fig. 4B-C, S4E).

To test if the StkyC domain was critical, we added back the StkyC domain (amino acids 167-224) to MBD6^CΔ^ (MBD6^CΔ+StkyC^). MBD6^CΔ+StkyC^ was able to rescue MBD6 nuclear foci counts, and complemented the derepression of *FWA* in the *mbd5 mbd6* mutant (Fig. 4B-E). Importantly, MBD6^CΔ+StkyC^ expressed in *acd15 acd21* mutant plants formed very few nuclear foci, similar to the low number of MBD6^CΔ^ foci in wild-type plants, demonstrating that foci localization rescue by the StkyC domain requires ACD15 and ACD21 (Fig. 4D).

To determine if the StkyC domain is responsible for localizing ACD15 and ACD21 to the MBD5/6 complex, we performed fluorescent protein colocalization experiments by co-expressing ACD21-CFP or ACD15-YFP with MBD6-RFP, MBD6^CΔ^-RFP, or MBD6^CΔ+StkyC^ RFP in *mbd5 mbd6* mutants. ACD15 and ACD21 both strongly correlated with full length MBD6 (Pearson correlation coefficient (*r)* of 0.96 and 0.86 respectively) and overlapped well with MBD6 signal across root nuclei, whereas ACD15 and ACD21 showed much weaker correlations with MBD6^CΔ^ (*r* = 0.67 and 0.46, respectively) and lost overlap with MBD6^CΔ^ nuclear signal (Fig. 4F-G and S4F-G). ACD15 and ACD21 also showed visibly higher cytosolic signal and lower nuclear signal when co-expressed with MBD6^CΔ^ in *mbd5 mbd6* (Fig. 4F-G and S4F-G). The addition of the StkyC domain (MBD6^CΔ+StkyC^) restored the correlation of ACD15 and ACD21 with MBD6 (*r* = 0.94 and 0.84 respectively), restored the overlap of ACD15 and ACD21 with MBD6 nuclear signal, and reversed the cytosolic localization of ACD15 and ACD21 (Fig. 4F-G and S3F-G).

To further test if ACD15 is needed for ACD21 to associate with MBD6^CΔ+StkyC^, we colocalized ACD15 and ACD21 with MBD6^CΔ+StkyC^ in wild-type or *acd15 acd21* double mutant plants (Fig. 4H). ACD21 showed a reduced correlation with MBD6^CΔ+StkyC^ in *acd15 acd21* plants compared to wild type (0.48 vs 0.79), a reduction of colocalization with MBD6 across nuclei, and a visible increase in ACD21 cytosolic localization, suggesting that ACD21 requires ACD15 to associate properly with MBD6^CΔ+StkyC^ (Fig.4I and S4H-I). On the other hand, ACD15 correlated strongly with MBD6^CΔ+StkyC^ in *acd15 acd21* plants (*r* = 0.92), maintained strong nuclear signal, and directly overlapped with nuclear MBD6, demonstrating that ACD15 does not require ACD21 for proper localization with MBD6^CΔ+StkyC^ (Fig. 4H and S4J). These experiments demonstrate that the StkyC domain of MBD6 is required for the function of the MBD5/6 complex, is needed for proper localization of ACD15 and ACD21, and mediates the accumulation of MBD6 at heterochromatic foci through ACD15 and ACD21.

### ACD15 and ACD21 can mediate functional and targeted gene silencing foci

Some ACD containing sHSP proteins are known to form dynamic oligomeric assemblies as part of their function in maintaining protein homeostasis (*16*), which could explain how ACD15/ACD21 drive high levels of MBD5/6 complex accumulation at meCG dense heterochromatin. To further explore this concept, we created a system to target MBD5/6 complexes to a discrete genomic location outside of pericentromeric heterochromatin. We utilized the SunTag system(*38*), composed of a dead Cas9 protein (dCas9) fused to ten single-chain variable fragment (scFv) binding sites, targeted to the promoter of the euchromatic *FWA* gene (*39*). To nucleate MBD5/6 foci at the dCas9 binding site we fused the scFv to the StkyC domain of MBD6 and to GFP, to visualize the nuclear distribution of the fusion proteins (Fig. 5A).

**Fig. 5.**
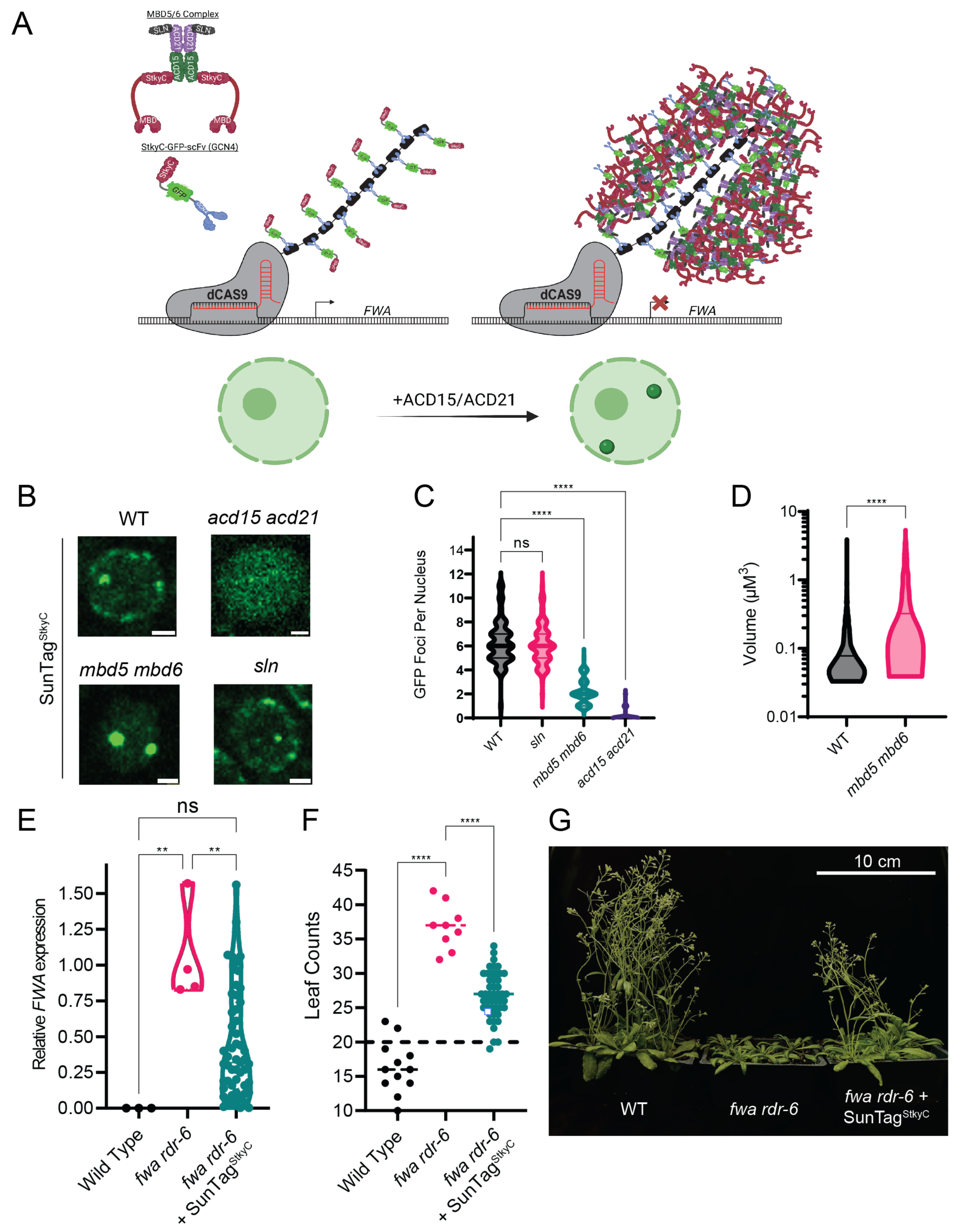
ACD15 and ACD21 drive the formation of MBD5/6 multimeric assemblies. **(A)** Graphical representation of SunTag^StkyC^ system and the hypothesized result. Created with BioRender.com. **(B)** Representative nuclear images of SunTag^StkyC^ in different mutant backgrounds. **(C)** SunTag^StkyC^ GFP foci counts per nucleus (N=100 per genotype). Compared using Welch’s ANOVA Dunnett’s T3 multiple comparisons test (****: P<0.0001, NS: P>=0.05). Scale bar = 2μM. **(D)** Volume of SunTag^StkyC^ GFP foci from 5 plant lines per genotype (WT: n=1461, *mbd5 mbd6*: n=1371). Two tailed t-test (****: P<0.0001, NS: P>=0.05). **(E)** RT-qPCR showing *FWA* expression in leaf tissue from T1 or control plants. Brown-Forsythe ANOVA with Tukey’s multiple corrections test (**: P<0.01, NS: P>=0.05). **(F)** Leaf counts post flowering of T1 *fwa rdr-6* SunTag^StkyC^ plants. Brown-Forsythe ANOVA with Dunnett’s multiple comparisons test (****: P<0.0001). **(G)** Representative image of early flowering T2 *fwa rdr-6* plants expressing SunTag^StkyC^.

If ACD15 and ACD21 drive higher order multimerization of MBD5/6 complexes, we would expect to observe discrete GFP foci in nuclei representing the dCas9 binding sites, as well as other foci corresponding to chromocenters since the scFv-GFP-StykC fusion would likely be recruited into multimerized MBD5/6 complexes at heterochromatin sites (Fig. 5A). Indeed, we observed an average of 6.4 GFP foci per nucleus in SunTag^StkyC^ expressing wild-type plants (Fig. 5B, C), some of which overlapped with DAPI staining chromocenters and others that did not (fig. S5A). We also transformed SunTag^StkyC^ into the *mbd5 mbd6* mutant, which would be predicted to eliminate recruitment of the scFv-GFP-StykC fusion protein into chromocenters by elimination of meCG bound endogenous MBD5/6 complexes. As predicted, we now observed an average of only two foci per nucleus (Fig. 5B, C), likely corresponding to the *FWA* alleles on the two homologous chromosomes. Consistent with these foci representing dCas9 bound to euchromatic *FWA* (*39*), these foci did not overlap with DAPI staining chromocenters (fig. S5B). Notably, the volume of SunTag^StkyC^ foci were increased in *mbd5 mbd6* (Fig. 5D), with the vast majority of nuclear GFP signal accumulating at the two nuclear bodies (Fig. 5B), suggesting that excess scFv-GFP-StykC fusion protein shifted from heterochromatic regions to the dCas9 binding sites.

We also expressed the SunTag^StkyC^ system in *acd15 acd21* mutants to determine if ACD15 and ACD21 are required for foci formation. Indeed, SunTag^StkyC^ now only displayed diffuse nucleoplasmic GFP signal, lacking detectable foci (Fig. 5B, C). This pattern was similar to control plants expressing a SunTag-TET1 system(*40*), in which the scFv was fused to GFP and the human TET1 protein, suggesting that the GFP foci observed in SunTag^StkyC^ is not a general property or artifact of the SunTag system (fig. S5C). We also introduced the SunTag^StkyC^ system into the *sln* genetic background and observed GFP foci counts and localization similar to the wild-type plants, showing around 6.1 foci per nucleus (Fig. 5B,C). These results demonstrate that ACD15/21 are necessary and sufficient to drive high level accumulation of MBD5/6 complexes at discrete foci.

We next tested if the foci formed by the SunTag^StkyC^ system are capable of gene silencing. The *FWA* gene is normally methylated and silent in wild-type plants. However, stably unmethylated and expressed *fwa* epigenetic alleles exist that cause a later flowering phenotype (*41, 42*). This allowed us to test if the SunTag^StkyC^ system could silence *FWA* by introducing the system into the *fwa* epigenetic background. Indeed, we found a significant suppression of *FWA* expression compared to *fwa* control plants (Fig. 5E). *fwa* plants expressing SunTag^StkyC^ also flowered earlier on average, showing a decrease in the number of leaves produced before flowering compared to *fwa* mutant plants (Fig. 5F-G). Correlation of *fwa* expression with leaf counts for *fwa* SunTag^StkyC^ plants showed a strong positive correlation as expected (fig. S5D).

Lastly, we tested whether the SunTag^StkyC^ system could complement the *FWA* derepression phenotype of MBD5/6 complex mutants (Fig. 1E)(*29*). Interestingly, SunTag^StkyC^ was able to silence *FWA* in *mbd5 mbd6* mutant plants demonstrating that the tethering function of MBD6 could be largely replaced by targeting with the StkyC domain, and that silencing can occur without the methyl binding proteins (fig. S5E). Surprisingly, SunTag^StkyC^ could also partially complement *FWA* derepression in the *sln* mutant background, while SunTag^StkyC^ could not complement *FWA* derepression in the *acd15 acd21* mutant background (fig. S5F,G). These results demonstrate that the SunTag^StkyC^ system maintains some gene silencing capability without SLN, suggesting that ACD15 and ACD21 alone possess silencing ability.

### Concluding remarks

Our results provide evidence for distinct mechanistic roles for ACD15, ACD21, and SLN in the formation and regulation of the meCG specific MBD5/6 silencing complex (fig. S6). ACD15 and ACD21 function to both drive the formation of higher order MBD5/6 complex assemblies, and bridge SLN to the complex. In contrast, the main role of SLN appears to be regulation of the dynamics of protein mobility within these complex assemblies. The activity of both ACD15/21 and SLN are clearly required for proper silencing function of the complex. The accumulation of multiple MBD5/6 proteins into higher order complex assemblies can explain why these complexes preferentially localize to high density meCG sites in the genome, likely via cooperative binding to closely spaced meCG sites.

ACD domain containing small HSPs are found in all eukaryotic lineages and are most well known for their role in regulating the aggregation of proteins (*14, 15, 17, 34, 43*). In the MBD5/6 complex however, the oligomerization capacities of ACD15 and ACD21 are specifically co-opted to control complex multimerization and silencing function. It seems likely that ACD proteins in other systems may also play important roles outside of general protein homeostasis.

## Acknowledgments

We thank Suhua Feng, Mahnaz Akhavan and the Broad Stem Cell Research Center Biosequencing core for DNA sequencing, and Julie Law for *lil-1* mutant seeds.

## Funding

National Institutes of Health grant R35 GM130272 (SEJ), R01 GM089778 (JAW)

The Philip Whitcome Pre-Doctoral Fellowship in Molecular Biology to B.A.B and to L.I. SEJ is an Investigator of the Howard Hughes Medical Institute

## Author contributions

Conceptualization: BAB, LI, SEJ

Investigation: BAB, LI, SW, JY, YJA, JS, CM, BS, AVA, SKL, IT

Resources: SEJ Software: BAB, LI, SW

Validation: BAB, LI, SW

Data curation: BAB, LI

Formal Analysis: BAB, LI, SW

Funding acquisition: BAB, LI, SEJ

Methodology: BAB

Visualization: BAB, LI, SW

Project administration: BAB, SEJ

Supervision: JAW, SEJ

Writing –original draft: BAB, LI, SEJ

Writing –review & editing: BAB, LI, SEJ

## Competing interests

Authors declare that they have no competing interests.

## Data and materials availability

The high-throughput sequencing data generated in this paper have been deposited in the Gene Expression Omnibus (GEO) database (accession no. TBD).

## Supplementary Materials

Materials and Methods

Supplementary Text

Figs. S1 to S6

Table S1

References (*1*–*57*)

## Notes

### Competing Interest Statement

The authors have declared no competing interest.

### Summary of Updates

There are few major deferences between the revised version and the previous version of the manuscript

