## Supplemental Materials for "ACD15, ACD21 and SLN regulate accumulation and mobility of MBD6 to silence genes and transposable elements"

**The PDF file includes:**

Materials and Methods
Figs. S1 to S6

**Other Supplementary Materials for this manuscript include the following:**

Table S1

**Materials and Methods**

Plant materials and growth conditions

All plants used in this study were in the Columbia-0 ecotype (Col-0) and were grown on soil in a greenhouse under long-day conditions (16h light / 8h dark). Plants grown for microscopy were plated on 1/2MS plates in growth rooms at room temperature (~25°C), with 16h of light and 8h of dark.

The following mutant lines were previously described: *mbd5 mbd6* T-DNA double mutant composed of *mbd5* T-DNA line SAILseq_750_A09.1 and *mbd6* T-DNA line SALK_043927 (*29)*; *mbd5 mbd6* double mutant composed of *mbd5* CRISPR/Cas9-generated indel and *mbd6* T-DNA mutation SALK_043927 (*29)*; *sln* (SALK_090484) (*29*), *fwa rdr6-15 (41*), *lil-1 (30).* Novel mutants and transgenic lines were generated as described below.

Generation of CRISPR lines

CRISPR/Cas9 mutants for ACD15.5 and ACD21.4 were generated with the pYAO::hSpCas9 system (*44)*. We designed two guide RNAs per gene with the goal of generating large deletions, one of them targeting the beginning of the coding region and another one targeting the end of the gene (Figure S1B). We were not able to obtain large deletions at these loci, but we found small indels causing frameshifts (Figure S1B). The guide RNAs were cloned sequentially in the AtU6-26-sgRNA cassette by overlapping PCR. The PCR product was cloned into the SpeI site of the pYAO::hSpCas9 destination plasmid by In-Fusion (Takara, 639650). The procedure was repeated four times (two guides for each gene). The final vector was electroporated into AGLO agrobacteria and transformed in Col0 or *sln* mutant plants (SALK_090484). T1 plants were selected on ½ MS agar plates with hygromycin B and were genotyped by PCR and by sanger sequencing of PCR amplified genomic regions surrounding each guide RNA. The lines containing the desired mutations were propagated to identify null segregants for the Cas9 transgene, and to obtain homozygous mutations. Experiments were performed in T4 generation.

Generation of transgenic lines

The transgenic lines expressing FLAG-tagged constructs used for IP-MS and ChIP-seq were generated as follows. Genomic DNA was cloned into pENTR/D-TOPO vectors (Thermo Fisher), including endogenous promoters and introns, until the last base before the STOP codon. The MBD5 gene was cloned starting from 1094 bp before the TSS, MBD6 from 294 bp before the TSS, SLN from 2351 bp before the TSS, ACD15.5 from 644 bp before the TSS, and ACD21.4 from 266 bp before the TSS. The genes were then transferred via a Gateway LR Clonase II reaction (Invitrogen, 11791020) into a pEG302 based binary destination vector including a C-terminal 3xFLAG epitope tag. The final vectors were electroporated into AGL0 agrobacteria that were used for plant transformation by agrobacterium-mediated floral dipping. T1 transgenic plants were selected with hygromycin B on ½ MS agar medium or with Basta (Glufosinate) on soil. IP-MS and ChIP-seq experiments were done in T2 or T3 generation.

Transgenic plants expressing fluorescently tagged proteins were created using the pGWB553 (https://www.addgene.org/74883/), pGWB540 (https://www.addgene.org/74874/), and pGWB543 (https://www.addgene.org/74877/). Specifically, ACD15, ACD21, and SLN promoters and coding sequences were PCR amplified from genomic DNA (as explained above) and cloned into pENTR vectors. These coding sequences were then inserted into final destination vectors using Gateway LR Clonase II Enzyme mix(Catalog number: 11791020, ThermoFisher). These final destination vectors were then electroporated into AGLO and transformed into Col0, *mbd5 mbd6* (SALK_043927), *sln (*SALK_090484*), acd15 acd21*, *acd21*, *acd15*, and *acd15 acd21 sln* (SALK_090484) mutant plants. Positive T1 plants were selected on ½ MS agar plates with hygromycin B and confirmed by western blots using fluorescent protein specific antibodies.

Transgenic plants expressing $\text{SunTag}^{\text{StkyC}}$ were created from a previously published $\text{SunTag}^{\text{TET1}}$ plasmid using the StkyC domain sequence (amino acids 173-225) of MBD6 (*45)*. The $\text{SunTag}^{\text{StkyC}}$ was targeted using two guides (Guide 4 (ACGGAAAGATGTATGGGCTT) and Guide 17 (AAAACTAGGCCATCCATGGA)) which were cloned as previously described (*46*). This plasmid was electroporated into AGLO and transformed into Col0, *mbd5 mbd6* (SALK_043927), *acd15 acd21*, *acd21*, *acd15*, and *sln* (SALK_090484), and *fwa rdr-6-15(41).* Positive selection of transgenic plants was done on ½ MS agar plates with hygromycin B after 5 days in the dark at 4C, 8 hours in the light at room temperature, and another 5 days in the dark at room temperature.

<u>Immunoprecipitation-Mass Spectrometry</u>
IP-MS experiments were performed as previously described (*29)*. Briefly, 8 to 10 g of inflorescences for each sample were used. Frozen tissue was ground with a tissue lyser and resuspended in IP buffer (50 mM Tris·HCl pH 8.0, 150 mM NaCl, 5 mM EDTA, 20% glycerol, 0.1% Tergitol, 0.5 mM DTT, and cOmplete EDTA-free Protease Inhibitor Cocktail [Roche]). Samples were filtered with miracloth, disrupted with a Dounce homogenizer, and centrifuged for 10 min at 4°C at 20,000 g. Supernatants were incubated with 200 µL of M2 magnetic FLAG-beads (SIGMA, M8823) for 2 hours rotating at 4°C. The beads were washed 5 times in IP buffer and eluted with 250 µg/mL 3X-FLAG peptides in TE. The eluted protein complexes were precipitated overnight with 20% trichloroacetic acid (TCA).
<u>Digestion and Desalting</u>
The protein pellets were resuspended with 100 µl digestion buffer (8M Urea, 0.1M Tris-HCl pH 8.5). Then the samples were reduced and alkylated via sequential 20-minute incubations with 5 mM TCEP and 10 mM iodoacetamide at room temperature in the dark while being mixed at 1200 rpm in an Eppendorf thermomixer. 20 µl of carboxylate-modified magnetic beads (CMMB, also widely known as SP3 (*47*) was added to each sample. Ethanol was added to a concentration of 50% to induce protein binding to CMMB. CMMB were washed 3 times with 80% ethanol and then resuspended with 50 µl 50 mM TEAB.
The protein was digested overnight with 0.1 µg LysC (Promega) and 0.8 µg trypsin (Thermo Scientific, 90057) at 37 °C. Following digestion, 1.2 ml of 100% acetonitrile was added to each sample to increase the final acetonitrile concentration to over 95% to induce peptide binding to CMMB. CMMB were then washed 3 times with 100% acetonitrile and the peptide was eluted with 65 µl of 2% DMSO. Eluted peptide samples were dried by vacuum centrifugation and reconstituted in 5% formic acid before analysis by LC-MS/MS.
<u>LC-MS Acquisition and Analysis</u>
Peptide samples were separated on a 75 µM ID, 25 cm C18 column packed with 1.9 µM C18 particles (Dr.Maisch GmbH) using a 140-minute gradient of increasing acetonitrile concentration ,and injected into a Thermo Orbitrap-Fusion Lumos Tribrid mass spectrometer. MS/MS spectra were acquired using Data Dependent Acquisition (DDA) mode.

MS/MS database searching was performed using MaxQuant (1.6.10.43) (*48*) against the Arabidopsis thaliana reference proteome TAIR (Araport11 release).

Chromatin Immunoprecipitation-sequencing (ChIP-seq)

The anti-FLAG ChIP-seq experiments were performed as previously described (*29)*. The RFP ChIP-seq experiments (Figure 3) were done with the following variations: 1) After sonication and two rounds of centrifugation, 50 µl of ChromoTek RFP-Trap Magnetic beads (proteintech, Cat No. rtma) were added to each sample for overnight incubation. 2) For elution, 250 µl elution buffer (SDS 1%, NaHCO3 0.1 M) was added, and samples were shaken for 15 min at room temperature. This step was repeated twice to reach 500ul of final elution volume. 480 µl of eluate was combined with 20 µl of 5M NaCl and incubated in a thermomixer overnight at 65C and 400rpm for reverse crosslinking. The following steps were performed as previously described (*29)*.

ChIP-seq libraries were prepared with the Ovation Ultra Low System V2 1-16 kit (NuGEN, 0344NB-A01) following the manufacturer's instructions, with 15 cycles of PCR. Final libraries were sequenced with the Illumina NovaSeq 6000 System.

ChIP-seq analysis

Raw reads were filtered based on quality score and trimmed to remove Illumina adapters using Trim Galore (Babraham Institute). Filtered reads were mapped to the Arabidopsis reference genome (TAIR10) with Bowtie2 (*49*) with default parameters. PCR duplicates were removed using MarkDuplicates.jar (picard-tools suite, Broad Institute). Genome browser tracks for visualization purposes were generated using deeptools (v 3.0.2) bamCoverage (*46*) with the options -- normalizeUsing RPKM and --binSize 10. To obtain tracks normalized over the no-FLAG control, we used deeptools bamCompare (*50*) with the "log2" option.
The analysis of correlation between ChIP-seq data and mCG density was performed as previously described (*29)*, by calculating the sum of CG methylation percentages in 400 bp bins. The data were plotted using the R package *ggplot* with the option *geom*_smooth.
ChIP-seq peaks were called with MACS2 (v 2.1.0) (*51*) using an FDR cutoff of 0.01. The FLAG and RFP associated hyperchippable regions, defined as peaks called in the anti-FLAG Col0 or anti-RFP Col0 controls, were subtracted from the peak sets of each sample. The peaks of individual replicates for ACD15 and ACD21 were merged with *homer mergePeaks* using the option -d given (*52*). Overlap analysis of different ChIP-seq peak sets was performed with *homer mergePeaks* using the options -d given and -venn (*52*).

RT-qPCR

RNA samples for RT-qPCR experiments were purified using Direct-zol RNA miniprep kit (catalog number: R2052, Zymo Research) from unopened flower bud tissue or leaf tissue used in Figure 5. cDNA samples were prepared using Superscript IV mastermix (catalog number: 11760500, Invitrogen) from ~400 ng of RNA and qPCR was performed using BioRad Sybergreen mastermix (catalog number: 1708882, Bio-Rad). Each qPCR experiment contained 2 technical replicates for each gene (either *FWA* or IPP2 housekeeping control). qPCR results were analyzed using BioRad CFX maestro software. *FWA* expression was normalized to expression of the reference gene *IPP2*, and to the control samples as indicated in each plot (i.e., *mbd5 mbd6*

mutants or *fwa rdr-6* mutant) using the ΔΔCq method. The data was graphed using GraphPad Prism software. Statistical analysis was done as described in the figure legends.
List of primers used for RT-qPCR:

| Primer name | Sequence |
|---|---|
| FWA RT-qPCR Fw | TTAGATCCAAAGGAGTATCAAAG |
| FWA RT-qPCR Rev | CTTTGGTACCAGCGGAGA |
| IPP2 RT-qPCR Fw | GTATGAGTTGCTTCTCCAGCAAAG |
| IPP2 RT-qPCR Rev | GAGGATGGCTGCAACAAGTGT |

RNA-Sequencing
RNA-sequencing was performed on mature pollen samples isolated as previously described *(53)*, with 6 biological replicates per genotype, grown and processed in 2 batches (3 replicates each). Briefly, 700-1000 µL of open flowers were harvested in 2-mL protein low bind tubes (Eppendorf). 700 µL of Galbraith buffer (45 mM MgCl2,30 mMC6H5Na3O7.2H2O [Trisodium citrate dihydrate], 20 mM MOPS, 0.1% [v/v] Triton X-100, pH 7) supplemented with 70 mM 2-Mercaptoethanol, were added to the tube, and the flowers were vortexed for 3 min at max speed in the cold room, to release the pollen from the anthers. The extraction procedure was repeated two times, and the two aliquots of pollen in solution were combined. The suspension was filtered with an 80 µm nylon mesh into a new 1.5 mL tube, and then spinned-down for 5 minutes at 500 g. The supernatant was carefully removed and the pollen was flesh frozen with a metal bead. Frozen samples were disrupted with a tissue grinder and RNA extraction was performed with the Zymo Direct-zol RNA MiniPrep kit (Zymo Research), with in-column DNase digestion. ~500 ng of RNA were used as input for library preparation using the TruSeq Stranded mRNA Library Prep Kit (Illumina), according to the manufacturer's instructions. The final libraries were sequenced with the Illumina NovaSeq 6000 System.

RNA-Sequencing analysis
RNA-sequencing reads were filtered based on quality score and trimmed to remove Illumina adapters using Trim Galore (Babraham Institute). The filtered reads were mapped to the Arabidopsis reference genome (TAIR10) using STAR (*54*) , allowing 5% of mismatches (-outFilterMismatchNoverReadLmax 0.05) and unique mapping (–outFilterMultimapNmax 1). MarkDuplicates from the Picard Tools suite was used to remove PCR duplicates. Coverage tracks for visualization in the genome browser were generated using Deeptools 3.0.2 bamCoverage with the options –normalizeUsing RPKM and –binSize 10 (*50*).
To obtain gene counts, we used a set of reference pollen transcriptome annotations that we previously generated (53*)* and are available from Github at https://github.com/clp90/mbd56_pollen. We used HTseq (*55*) with the option –mode = union, to obtain counts for all transcripts (genes, TEs, and other undefined non-coding transcripts). The HTseq gene counts were used to perform the differential gene expression analysis using the R package DEseq2 (*56*) with a cutoff for significance of adjusted p-value <0.05 and |log2FC|>1. Figures were generated using the R packages *ggplot* and *UpSetR*.
To determine the promoter CG methylation levels at each transcript (Figure 1D), we first identified promoters as a 600 bp region surrounding the TSS. Then we calculated average CG methylation percentages at promoters using bedtoolsmap (*57*) with the option ''mean''. Our previously published Col0 flower buds BS-seq dataset was used for this analysis and for the

representative genome browser tracks: GSM5026060 and GSM5026061 (combined replicates) (*29)*.

Amino Acid Alignment

Amino acid alignments of MBD5, MBD6, and MBD7 were performed using Clustal Omega multiple sequence alignment tool (https://www.ebi.ac.uk/Tools/msa/clustalo/). Amino acid sequences MBD5 (Accession No. Q9SNC0), MBD6 (Accession No. Q9LTJ1), MBD7 (Accession No. Q9FJF4) were obtained from UniProt protein database. The alignment was run with default settings.

AlphaFold Multimer Protein Structure Prediction

Protein structure predictions were run with the AlphaFold Colab notebook (AlphaFold.ipynb, https://colab.research.google.com/github/deepmind/alphafold/blob/main/notebooks/AlphaFold.ipynb#scrollTo=XUo6foMQxwS2 ) (*33).* The standard workflow was followed, and "run_relax" option was disabled. The .pdb output files were visualized with Pymol (Delano Scientific, LLC.).

Leaf Counting

Leaf counting was performed as mentioned previously (*29*) where total numbers of rosette and cauline leaves were counted in T1 generation of plants grown side-by-side under the same conditions.

Confocal Microscopy

All confocal microscopy experiments were performed using the LSM 980 confocal microscope. Unless otherwise stated, all experiments were performed using a 40x magnification water objective lens. For all experiments using multiple fluorescent tags, we manually gated the excitation and emission spectrum to limit any cross reactivity of the samples.

Live plant samples were prepared as follows:
2 week-old seedlings were grown on ½ MS plats at room temperature, ~25C, and then transferred using forceps onto 1mm thick glass slides (fisherscientific, Cat No.12-550-08) containing dionized water (room temperature). Seedlings were oriented such that root tips were on the middle of the slide while leaves were extending from the top of slides. #1.5 Coverslips (Fisherscientific, Cat No. 12-544-EP) were placed on top of the plant, gently, so as not to destroy or stress the seedling. Usually, 1-4 plants were placed on the one slide for imaging.

FRAP Experiment and Analysis

FRAP experiments were performed on a LSM 980 using 40x magnification and water objective lens. Images of a region of interested were obtained as a "snap" in order to circle a region of interest to be bleached. Then an experiment was run such that 5 images were taken followed by a bleaching event using 100% laser excitation wavelength, dependent on the fluorescent protein being imaged, for 300 iterations. Signal was then tracked post bleaching for indicated amount of time. FRAP analysis was performed using EasyFRAP online analysis software (https://easyfrap.vmnet.upatras.gr/). Briefly, three regions of interest were measured for each FRAP replicate: Bleached region (1), specific nucleus region containing the bleached foci (2), and a random region containing no signal in the root of the plant (3). The signal of these three regions across time were added into the given excel template from EasyFRAP and uploaded for analysis along with other replicate files (N=25 for each FRAP experiment). Plotted FRAP curves represent full normalization of the data to account for any variations in bleaching depth among samples. FRAP data starting from the bleaching event are plotted using GraphPad Prism software with 95% confidence intervals calculated from normalized FRAP data of FRAP experiment replicates. One-Phase association, non-linear regressions were fitted to estimate and statistically compare maximum plateau and $t_{1/2}$ for each FRAP experiment.

Quantification of foci counts, volume, and nuclear distributions.
All foci counts, volumes, and nuclear distribution plots were quantified using ImageJ, Image analysis software. Foci counts and volume measurements were obtained using 3D object counter from 50-slice z-stacks of root meristems across multiple plant lines using thresholding through ImageJ software. Nuclear distributions were obtained using plot profile feature across a fixed line length in ImageJ after converting images to RBG format. Intensities from nuclear distribution plots were then normalized to maximum intensity within each replicate to normalize the data distribution. Foci counts, volumes, and nuclear distribution intensity values were all plotted using GraphPad Prism software and statistical analysis was performed using GraphPad Prism software as mentioned in figure legends.

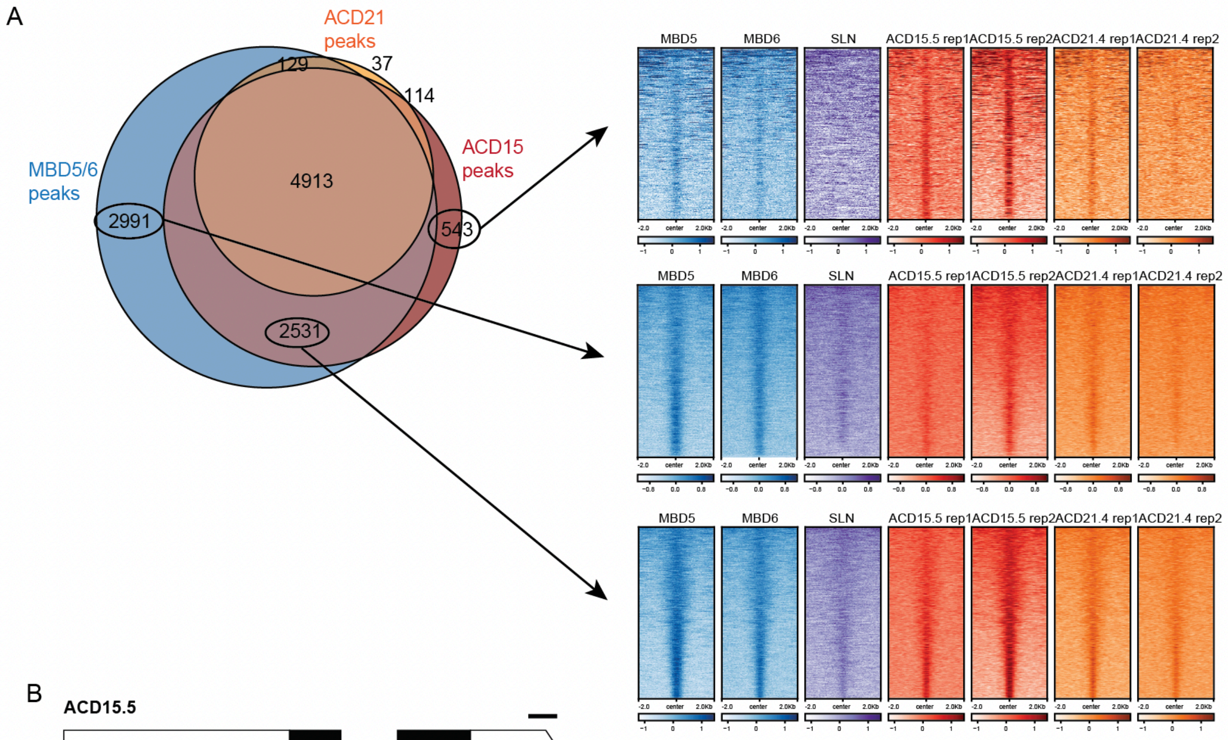


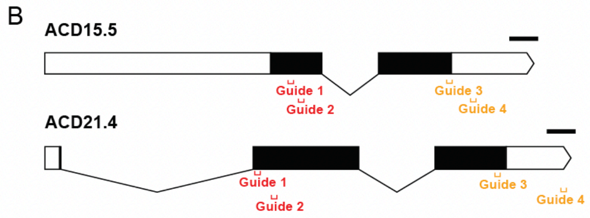


| | ACD15.5 | | ACD21.4 | | background |
|---|---|---|---|---|---|
| acd15.5 | guide 2 + 4 | Insertion at guide 2 causing STOP codon | guide 2 + 4 | WT | Col0 |
| acd21.4 | guide 1 + 3 | WT | guide 1 + 3 | single base insertion (A) at guide 1 | Col0 |
| acd15.5 acd21.4 | guide 1 + 3 | single base insertion (A) at guide 1 | guide 1 + 3 | 37bp deletion at guide 1 causing STOP codon | Col0 |
| adc15.5 acd21.4 sln | guide 1 + 3 | single base deletion (A) at guide 1 | guide 1 + 3 | single base insertion (A) at guide 1 | sln T-DNA: SALK_090484 |

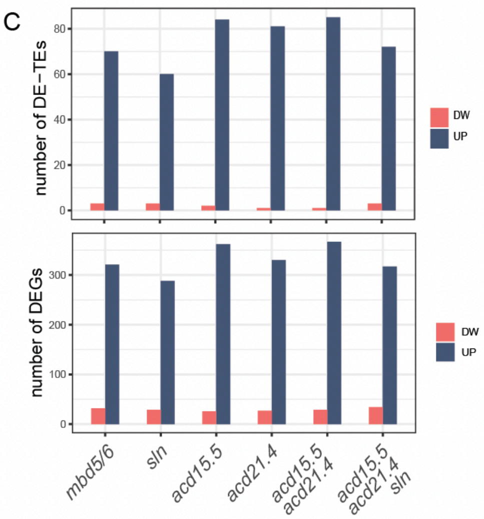


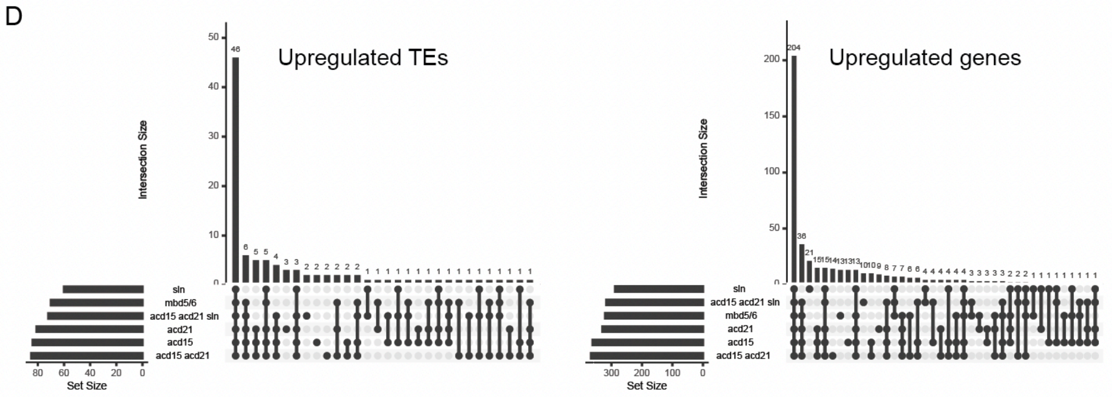

**Fig. S1. ChIP-seq and RNA-seq analysis of ACD15 and ACD21.**

**A)** Venn diagram of ChIP-seq peaks showing large overlap between samples. The peak sets indicated with circles (putative MBD5/6 unique, ACD15 unique, or MBD5/6/ACD15 unique peaks) were selected and visualized with heatmaps (right). We noted that at each peaks set groups, enrichment of most proteins was observed, thus suggesting that these regions are bound by all components of the MBD5/6 complex, despite not reaching our stringent significance threshold to be called as peaks. The heatmap shows log2(fold-change) over no-FLAG control. **B)** Scheme of ACD15 and ACD21 genes showing the location of the guide RNAs used for CRISPR/Cas9 mediated mutants generation. The table below shows the mutations obtained in each line. **C)** Barplots showing the number of differentially expressed TEs (DE-TEs) or differentially expressed genes (DEGs) in the indicated genotypes. **D)** Upset plots showing the intersection of the upregulated genes or TEs found for each genotype. The largest intersection group constitutes loci upregulated in all six mutant lines.

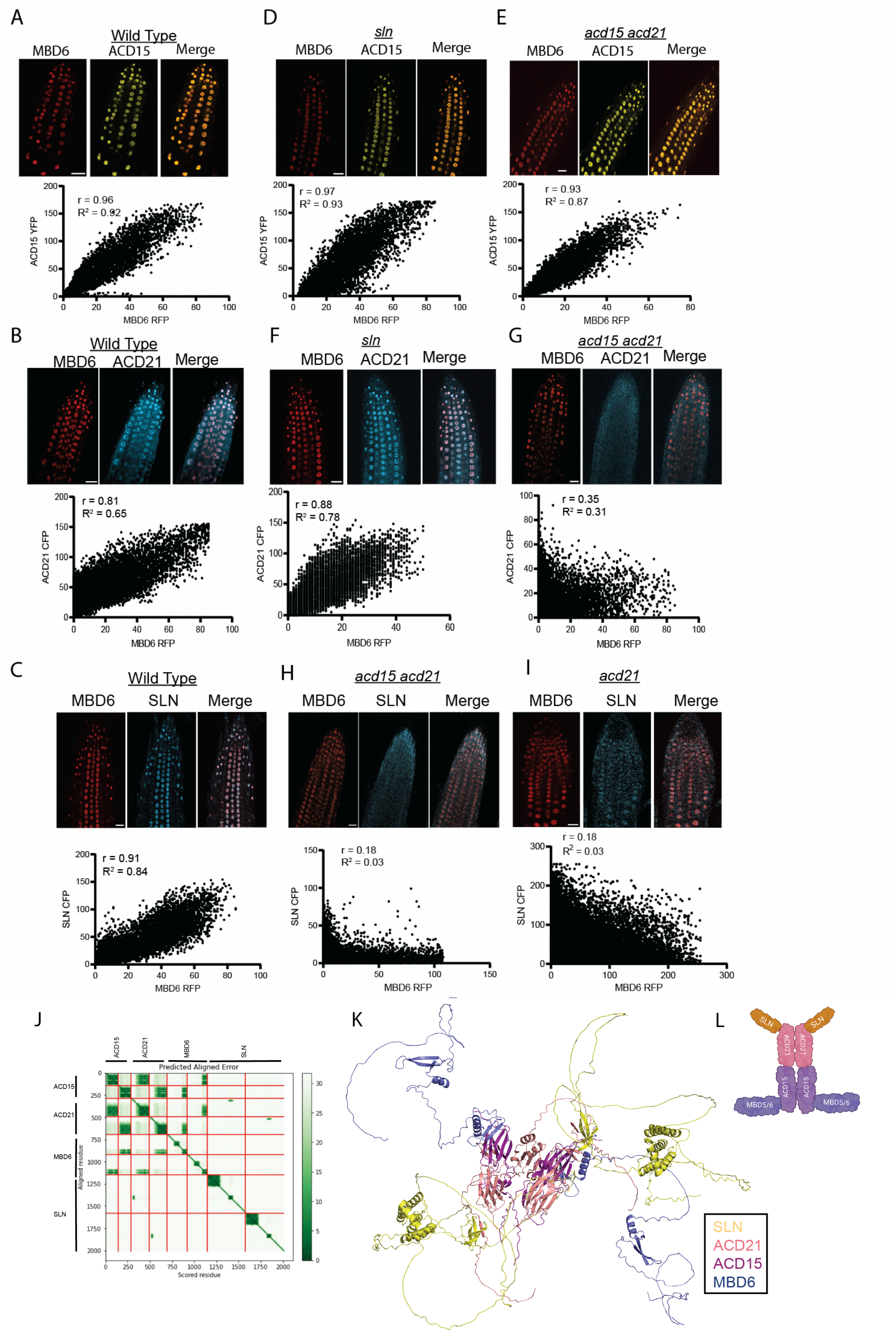
A
Wild Type
MBD6
ACD15
Merge
r = 0.96
R² = 0.92
ACD15 YFP
MBD6 RFP
D
sln
MBD6
ACD15
Merge
r = 0.97
R² = 0.93
ACD15 YFP
MBD6 RFP
E
acd15 acd21
MBD6
ACD15
Merge
r = 0.93
R² = 0.87
ACD15 YFP
MBD6 RFP
B
Wild Type
MBD6
ACD21
Merge
r = 0.81
R² = 0.65
ACD21 CFP
MBD6 RFP
F
sln
MBD6
ACD21
Merge
r = 0.88
R² = 0.78
ACD21 CFP
MBD6 RFP
G
acd15 acd21
MBD6
ACD21
Merge
r = 0.35
R² = 0.31
ACD21 CFP
MBD6 RFP
C
Wild Type
MBD6
SLN
Merge
r = 0.91
R² = 0.84
SLN CFP
MBD6 RFP
H
acd15 acd21
MBD6
SLN
Merge
r = 0.18
R² = 0.03
SLN CFP
MBD6 RFP
I
acd21
MBD6
SLN
Merge
r = 0.18
R² = 0.03
SLN CFP
MBD6 RFP
J
Predicted Aligned Error
ACD15
ACD21
MBD6
SLN
Aligned residue
Scored residue
K
SLN
ACD21
ACD15
MBD6
L
SLN
ACD21
ACD15
MBD5/6

**Fig. S2. Organization of the MBD5/6 complex structure.**
**(A-I)** Correlation between MBD6-RFP signal and either ACD15-YFP, ACD21-CFP, or SLN-CFP signal in the indicated mutant backgrounds (underlined). Images represent individual z-stack slices of roots from plants co-expressing MBD6 with either ACD15, ACD21, and SLN. Scatter plots indicate signal intensity for each fluorescent protein at each pixel of the image shown. Correlation coefficient: Pearson. Scale bars = 20 μM. **(J-K)** AlphaFold Multimer predicted structure of MBD5/6 complex with two copies each of MBD6, ACDC15, ACD21, and SLN along with confidence score map of the predicted complex. **(L)** Cartoon representation of the core dimeric MBD5/6 complex based on the AlphaFold Multimer prediction. The figure was created with Biorender.com.

A

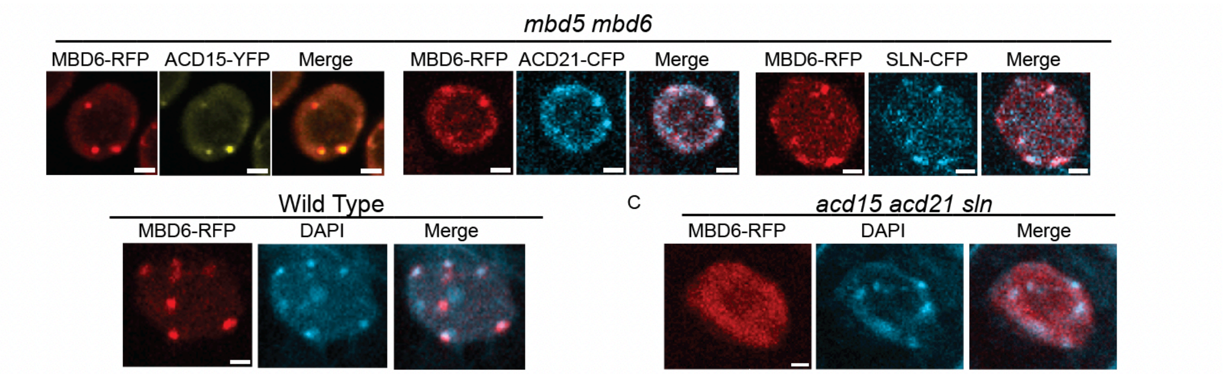


D

| | FRAP Recovery Half Time ($t_{1/2}$) | | Maximum Plateau Post Bleaching | |
|---|---|---|---|---|
| | **Wild Type** | ***sln*** | **Wild Type** | ***sln*** |
| MBD6 | 3.60 (3.175 to 3.991) | 2.44 (1.813 to 3.174) | 0.93 (0.9137 to 0.9561) | 0.61 (0.5826 to 0.6336) |
| ACD15 | 3.63 (3.243 to 4.046) | 4.93 (4.054 to 5.951) | 0.94 (0.9233 to 0.9635) | 0.59 (0.5676 to 0.6212) |
| ACD21 | 4.30 (3.735 to 4.927) | 3.26 (2.448 to 4.220) | 0.91 (0.8813 to 0.9351) | 0.63 (0.6009 to 0.6625) |

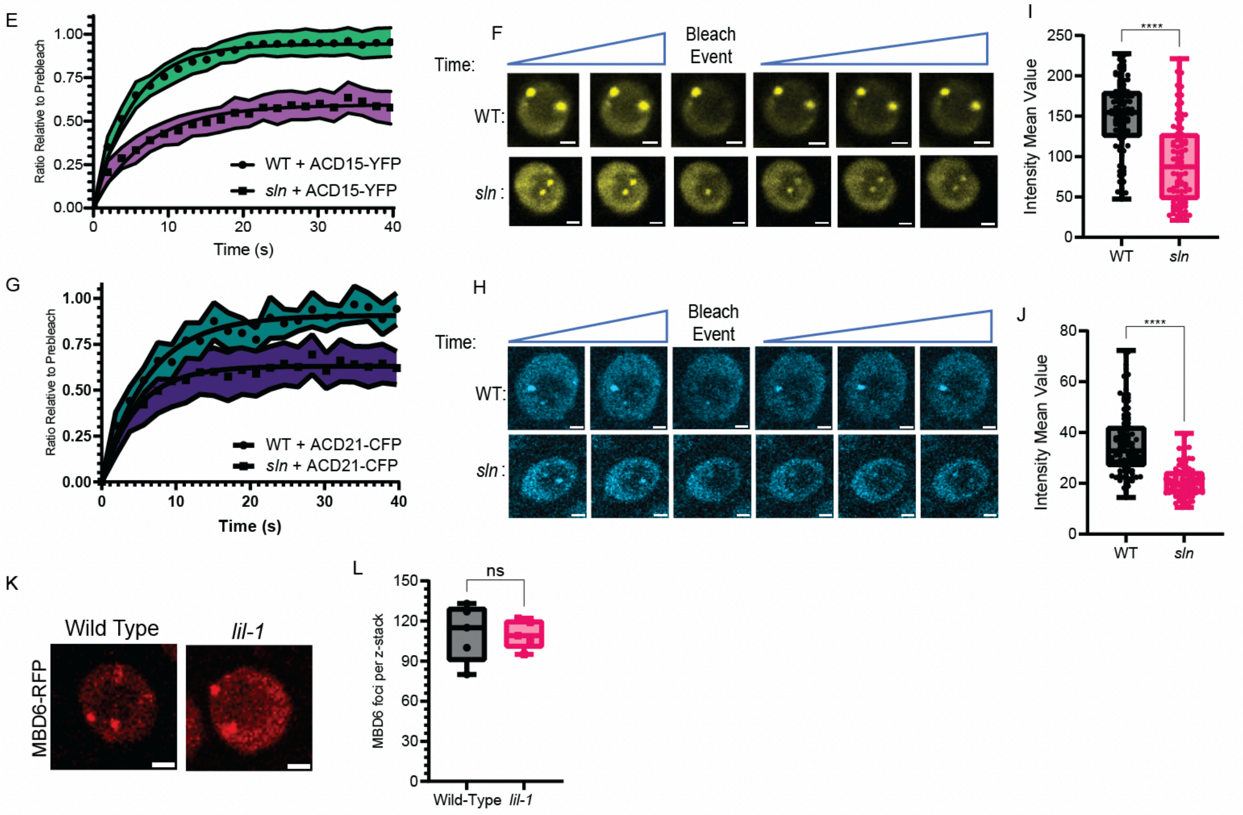


**Fig. S3. SLN regulates the nuclear mobility of MBD5/6 complex members.**

**(A)** Representative root nuclei images demonstrating MBD6-RFP overlap with ACD15-YFP, ACD21-CFP, and SLN-CFP. Scale bar = 2µM. **(B-C)** MBD6-RFP signal within DAPI-stained nuclei. Scale bar = 2µM. **(D)** Table of extrapolated values from FRAP curve data fitted with one-phase association liner regression using GraphPad Prism. **(E-H)** FRAP curves of ACD15 and ACD21 along with representative nuclei images of FRAP experiments. Shaded area: 95% confidence interval of FRAP data (N=25 from 5 plants lines), dots: mean values, line: fitted one-phase, non-linear regression. Scale bars = 2µM. **(I-J)** Intensity of ACD15 (I) and ACD21 (J) signal at 100 individual foci from multiple nuclei and plant lines. Comparisons were made using two-tailed t tests (****: P<0.0001). **(K)** Representative nuclei showing MBD6-RFP foci in lil-1 mutant and control plants. Scale bars = 2µM. **(L)** MBD6 foci counts across 50 slice Z-stacks of root meristem from five plant lines per genotype. Two Tailed T-test (NS: P>=0.05).

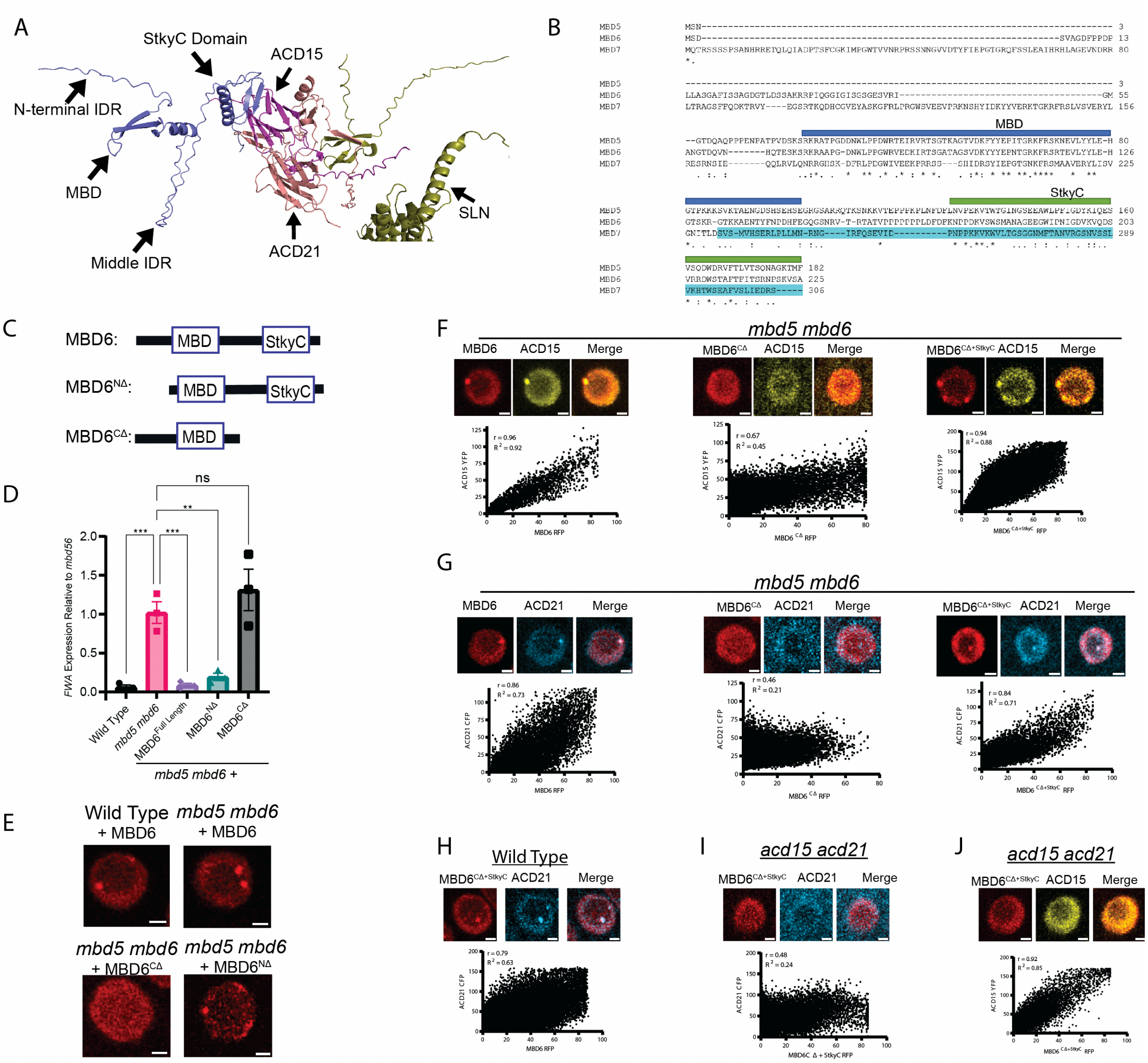

**Fig. S4. The StkyC domain is required for MBD6's function.**
**(A)** Protein structure representation of AlphaFold Multimer prediction of MBD6 with ACD15. Domains of MBD6 are annotated. **(B)** Protein alignment of MBD5, MBD6, and MBD7. MBDs and StkyC regions of MBD5, MBD6, and MBD7 are labelled. **(C)** Graphical representation of MBD6 deletion mutants. **(D-E)** RT-qPCR of *FWA* expression comparing *mbd5 mbd6* plants expressing MBD6 deletion mutants (D), along with representative images of MBD6 deletion mutants in root nuclei (E). Comparisons made using Brown-Forsythe ANOVA with Dunnett's multiple comparisons test. ***: P<0.001, **: P<0.01, NS: P>=0.05. Scale bars = 2μM. **(F-J)** Correlation of MBD6-RFP domain, deletion mutants with ACD15-YFP or ACD21-CFP along with representative nuclear images. Correlation coefficient: Pearson. Scale bars = 2μM.

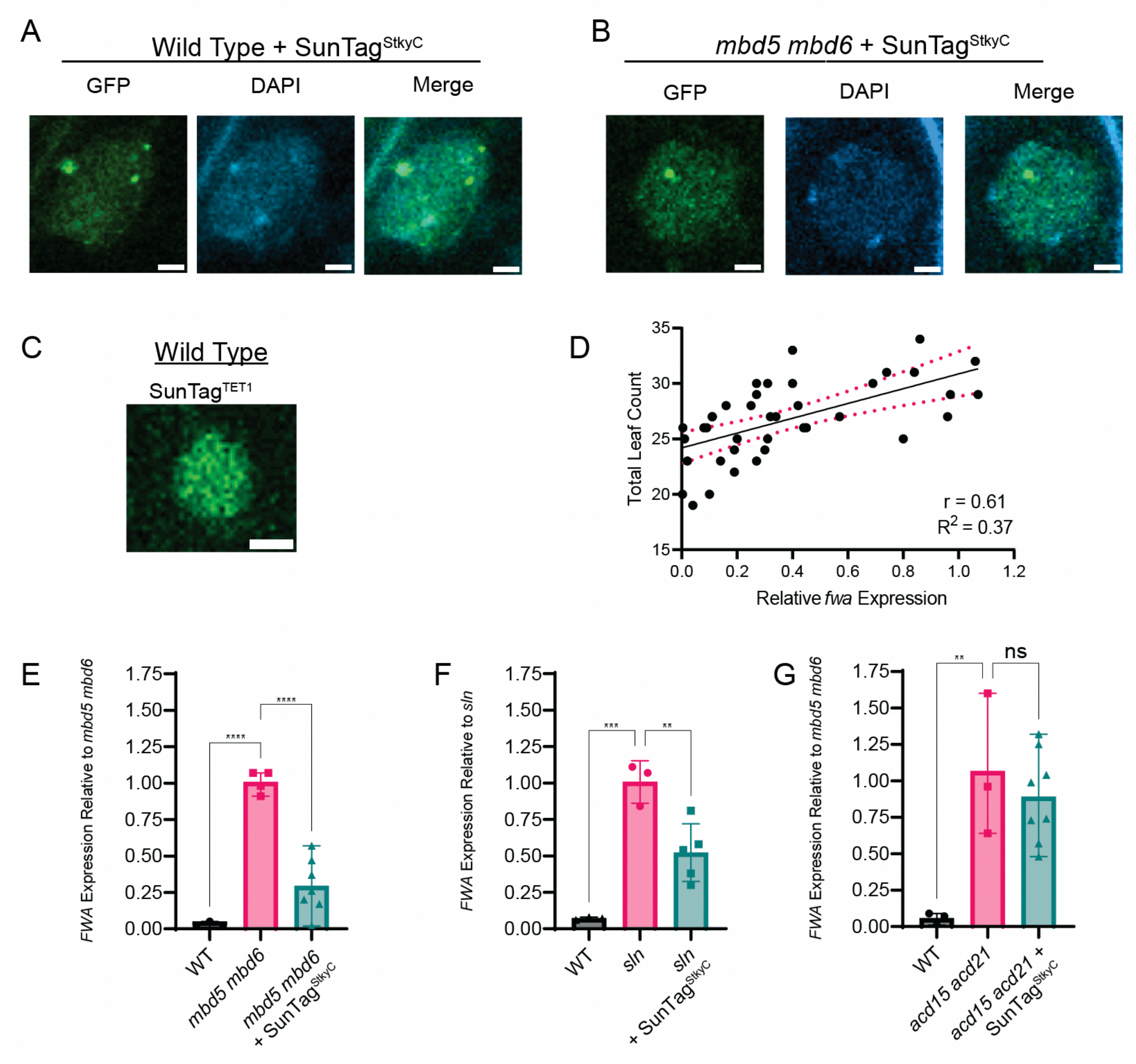


**Fig. S5. SunTag$^{StkyC}$ drives the formation of MBD5/6 nuclear foci.**
**(A-B)** SunTag$^{StkyC}$ expressing root nuclei stained with DAPI. Scale bars = 2µM. **(C)** Representative image of a root nucleus expressing SunTag$^{TET1}$. Scale bar = 2µM. **(D)** Correlation of SunTag$^{StkyC}$ *FWA* expression with leaf counts of individual T1 plants from Figure 5E-F. Correlation coefficient: Pearson. **(E-G)** RT-qPCR of *FWA* in *mbd5 mbd6, sln,* and *acd15 acd21* plants with and without SunTag$^{StkyC}$. Comparisons made using Brown-Forsythe ANOVA with Dunnett's multiple comparison test for each qPCR experiment. ****: P<0.0001, ***: P<0.001, **: P<0.01, *: P<0.05, NS: P>=0.05.

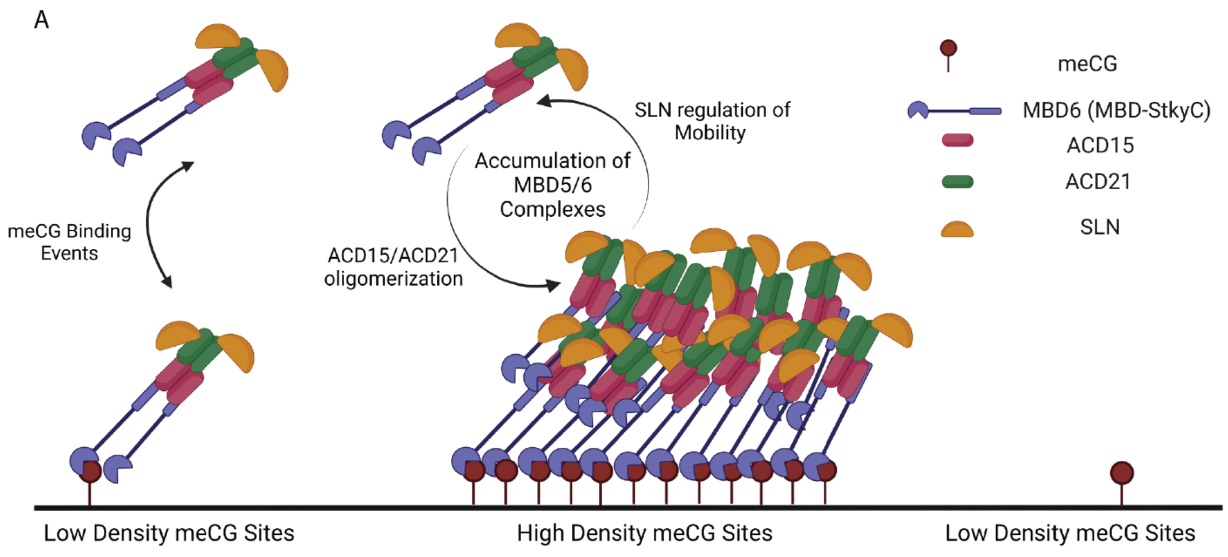


**Fig. S6. Model of MBD5/6 oligomerization at high density meCG sites.**
Diagram of proposed model showing ACD15/ACD21-dependent binding and accumulation of MBD5/6 complex members in multimeric assemblies. MBD5/6 recognize DNA methylation through their MBD domain. Although MBD5 or MBD6 can recognize individual meCG sites, regions with high density meCG sites facilitate recruitment of multiple MBD5/6 complexes, which triggers oligomerization: once MBD5/6 are bound to DNA, ACD15/ACD21 drive recruitment of other MBD5/6 complexes to facilitate oligomerization. This accumulation of proteins leads to higher than expected binding events and dwell time at meCG dense regions. SLN directly interacts with ACD21, accumulates with the complex, and acts to maintain the mobility of proteins within the oligomeric assembly. Created with BioRender.com.
